# An alternative model to prion fragmentation based on the detailed balance between PrP^Sc^ and suPrP

**DOI:** 10.1101/2020.04.24.058917

**Authors:** Monique Chyba, Jakob Kotas, Vincent Beringue, Christopher Eblen, Angelique Igel-Egalon, Yuliia Kravchenko, Human Rezaei

## Abstract

Prion assemblies responsible for transmissible spongiform encephalopathies grow in the form of linear amyloid fibrils. Traditional models for prion growth and tissue-spreading have relied upon the assumption that propagation is based on the process of fragmentation, wherein an assembly literally breaks in two, creating additional templating interfaces. Recent experimental data shows that PrP^Sc^ assemblies are in detailed balance with an elementary oligomeric building block called suPrP. In the present work we compare the dynamics of the canonical model of induced-fragmentation to the model of PrP^Sc^ assemblies in detailed balance with suPrP. The model is a dynamical system describing the populations of fibrils of varying lengths as a function of time; we analyze the system via both analytical and numerical techniques. We demonstrate that the detailed balance between suPrP and PrP^Sc^ model can equivalently replace the induced-fragmentation model. This equivalence opens a new opportunity in an optimal control problem.

## 1.1 Introduction

Transmissible spongiform encephalopathies (TSEs) are a group of rare and invariably fatal neurodegenerative diseases. Well-known examples include bovine spongiform encephalopathy (BSE) in cattle, scrapie in sheep, and Creutzfeldt-Jakob disease (CJD), Gerstmann-Sträussler-Scheinker syndrome (GSS), and fatal familial insomnia (FFI) in humans. A hallmark of TSEs is the conformational change of normal monomeric prion protein PrP^C^ into the abnormal aggregated assemblies called PrP^Sc^. The infectious process takes place via molecular templating, where the template molecule (PrP^Sc^) provides a pattern for the de novo generation of PrP^Sc^ assemblies. This simplest replication model constitutes the bedrock of the prion paradigm and defines the catalytic template assisted conversion of host-encoded monomeric PrP^C^ protein into misfolded assemblies PrP^Sc^. The prion paradigm is now extended to a number of neurodegenerative proteinopathies such as Alzheimer’s and Parkinson’s [7]. Despite important breakthroughs, the current paradigm fails to address the molecular mechanisms of prion replication due to the absence of PrP^Sc^ atomic structure and the coupling between linear-templating, leading to PrP^Sc^ assemblies’ size increase and prion amplification due to the multiplication of templating interfaces (Figure 1.1). The process of templating interface multiplication is central in the prion paradigm and is still not fully understood.

**Fig. 1.1.**
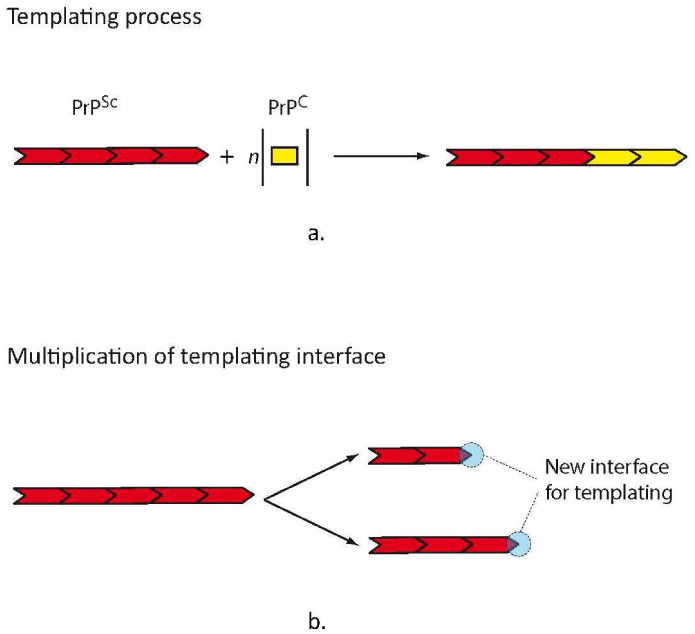
**a.** Process of prion replication by templating process leading to the size increase of PrP^Sc^ assemblies. The elongation by templating keeps the number of templating interfaces constant. **b.** The fibril fragmentation process has been proposed to explain the multiplication of templating interfaces and therefore prion amplification.

The role of heat shock proteins (Hsp) in fungus prion fragmentation revealed first by Kryndushkin and collaborators [12] and later by Shorter and Linquiest [16] brings a molecular explanation for the multiplication of templating interfaces correlated with the linear-growth of PrP^Sc^ assemblies’ size. However, the implication of Hsp in the fragmentation has been established for fungus prions. The fragmentation induced by Hsp machinery constitutes an active fragmentation requiring energy consumption provided by ATP or GTP hydrolysis [15, 16, 17]. The replication media of fungus prion intrinsically differs from those of mammalian prion. Indeed, mammalian prion replicates in an extracellular environment devoid of Hsp machinery and energetic sources when fungus prions are intracytoplasmic in the vicinity of housekeeping proteins, Hsp and sources of energy. Therefore, the question of how the multiplication of templating interfaces found at the source of prion replication and spreading occurs in mammalian prions is raised. Recent work demonstrated the existence of a constitutional detailed balance between PrP^Sc^ assemblies and an oligomeric conformer called suPrP. The latter constitutes the elementary building block of PrP^Sc^ assemblies [9, 10](Figure 1.2a). The existence of a detailed balance between PrP^Sc^ and suPrP makes the quaternary structure of prion assemblies highly dynamic in contrast with the widespread deadpan vision of amyloid assemblies. The suPrP harbors all the prion pathological determinant [10] and due to its small size, estimated between dimer or trimer, it could be involved in the spreading of the replicative center by simple diffusion rather than large PrP^Sc^ assemblies. According to the detailed balance process between PrP^Sc^ and suPrP, during prion replication the consumption of monomeric PrP^C^ also contributes to the increase of suPrP quantity (Figure 1.2b). This suPrP leakage could circumvent the necessity of fragmentation to amplify the templating interface [9]. Therefore, in the present work we explore the existence of the detailed balance between PrP^Sc^ and suPrP as an alternative process to the fragmentation to couple the linear templating process and templating interface amplification. Through theoretical kinetic analysis we demonstrated that linear increase of PrP^Sc^ size by templating is correlated with a non-linear increase in the amount of suPrP.

**Fig. 1.2.**
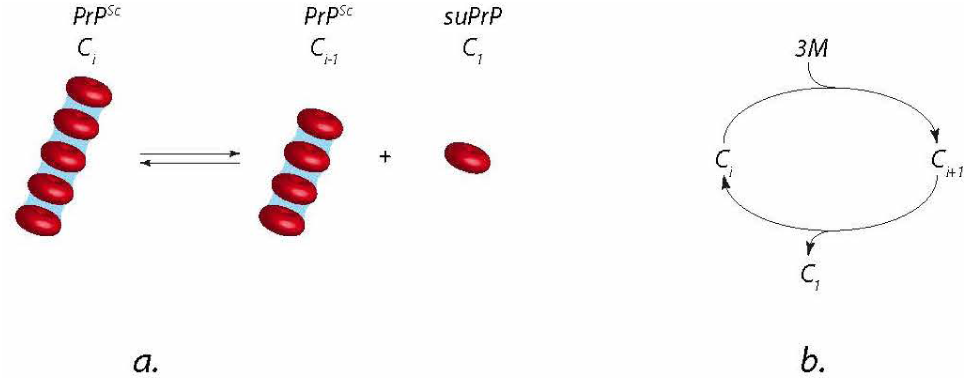
**a.** Existence of detailed balance between PrP^Sc^ assemblies and their elementary building block suPrP. **b.** The elementary kinetic scheme linking the detailed balance between PrP^Sc^ and suPrP and the templating process converting PrP^C^ to PrP^Sc^. *C*_*i*_ and *C*_*i*+1_ correspond to PrP^Sc^ assemblies of size *i* and *i* + 1 with *i* representing the number of suPrP.

In [14], a model of nucleated polymerization is introduced and in [3, 4] the authors consider a finite version of this model on which they apply optimal control to simulate the protein misfolded cyclic amplification (PMCA) protocol [5]. Fragmentation was the key process in this work to simulate prion replication. While we introduce a new model in this paper based on elementary building blocks to replace fragmentation, the same approach can be applied. In particular our simulations will show that the same behavior occurs in the evolution of the polymer density function. The outline of this paper is as follows. We begin by reviewing a widely studied dynamical system model where fragmentation is assumed to be the mechanism of prion replication. We use analytical techniques to find equilibrium points of the system; we then perform numerical simulations under assumptions on the parameter values. Finally, we perform sensitivity analysis on those parameters to understand their qualitative effect on the system. We then move to our novel suPrP model, also a dynamical system, where fragmentation has been replaced with the notion of oligomeric blocks. After building up the model, we again find equilibrium points and perform numerical simulations to understand the dynamics of the system for various parameter combinations. Finally, the paper introduces an optimal control problem by introducing prion assemblies evolving dynamically as a function of time and temperature. A three dimensional version of the system is analyzed using the tool of geometric optimal control.

### 1.2 Fragmentation Model

We begin by reviewing the widely studied model proposed by Masel, Jansen and Nowak [14], in which nucleated polymerization is assumed as the mechanism of prion replication. In this model, PrP^Sc^ are infectious agents and are linear on the macroscopic scale. The PrP^C^ are non-infectious monomers which form the building blocks of PrP^Sc^.

#### Notation

Let *x* denote the concentration of PrP^C^ monomers and *y*_*i*_ denote the concentration of PrP^Sc^ polymers of length *i*. We define *y* = ∑_*i*_ *y*_*i*_ to be the total concentration of PrP^Sc^ polymers, and introduce *z* = ∑_*i*_ *iy*_*i*_ as the total concentration of PrP^Sc^ subunits.

The model is a set of coupled differential equations incorporating the rates of the various relevant biological processes. Monomers are produced at the constant rate *λ* and degrade at a rate proportional to their concentration *x* with constant of proportionality *d*. Polymers of length *i* degrade at a rate proportional to their concentration *y*_*i*_ with constant of proportionality *a*. It is assumed that monomers degrade much more easily than polymers and thus *a* ≪ *d*.

#### Assumption: Polymerization and Fragmentation

This model assumes that monomers attach directly to a polymer of length *i* at a rate proportional to the product of their concentrations with constant of proportionality *β*. Moreover, polymers of length *i* ≥ *n* fragment into two pieces of size *j* and *i* − *j* at a rate proportional to their concentration with constant of proportionality *b*, where *n* is the critical size below which polymers are unstable and instantly disintegrate into PrP^C^ (thus *y*_*i*_ = 0 for all *i* < *n*).

It follows that monomer concentration decreases at a rate *βxy*. The concentration of polymers of length *i* increases at a rate *βxy*_*i*−1_ due to polymers of length *i* − 1 becoming polymers of length *i*, while simultaneously decreasing at a rate *βxy*_*i*_ due to polymers of length *i* becoming polymers of length *i* + 1. This leaves us with a net rate of change *βx*(*y*_*i*−1_ − *y*_*i*_) for the concentration of polymers of length *i*. The disease progresses slowly as the polymer chains fragment into smaller ones, thus creating more infectious assemblies onto which PrP^C^ can attach, effectively spreading the disease. Figure 1.3 depicts the assumptions made for this nucleated polymerization model.

**Fig. 1.3.**
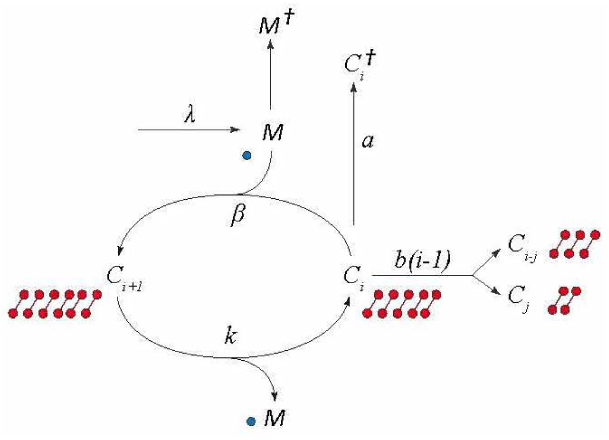
Canonical kinetic model of prion replication including fragmentation

This process is described by the following equations:

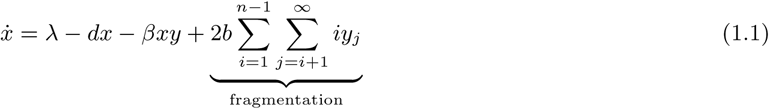

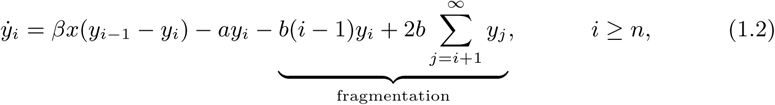

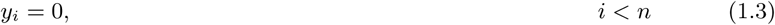

where *y* = ∑ *y*_*i*_ and the terms indicated with under-brackets are unique to the fragmentation process. Note that the *dx* term appearing in equation (1.1) should be interpreted as the rate constant *d* multiplied by the monomer population *x*, not as an infinitesimal.

Masel et al. show in their paper [14] that equations (1.1) - (1.3) can be closed under addition of the index *i*, resulting in a low-dimensional system:

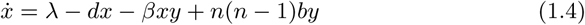

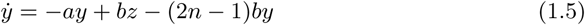

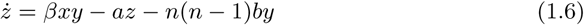

where *z* = ∑*iy*_*i*_ is the assembly concentration. There are two cases to distinguish: *b* = 0 and *b* ≠ 0.

#### Case *b* = 0

This corresponds to the situation with no fragmentation; that is, no depolymerization. It can be shown that equations (1.4) - (1.6) have a single equilibrium point at

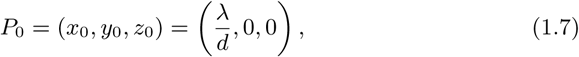

and that the Jacobian of equations (1.4) - (1.6) evaluated at *P*_0_ is

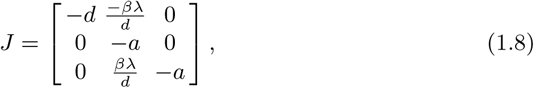

whose eigenvalues are −*d* and −*a* (with multiplicity two). Since *a* > 0 and *d* > 0, all eigenvalues are real and negative; thus it follows that equilibrium point *P*_0_ is asymptotically stable.

#### Case *b* ≠ 0

When fragmentation occurs, the situation is more complex. Equations (1.4) - (1.6) have two equilibrium points, one at *P*_0_ identical to the case *b* = 0, and one given by *P*_1_ = (*x*_1_, *y*_1_, *z*_1_), where

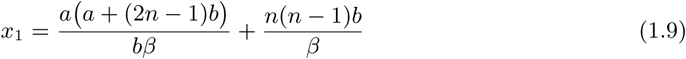

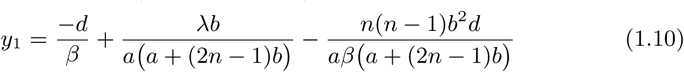

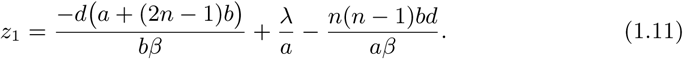

Note that lim_*b*→0_ *z*_1_ = −∞; also note that *z*_1_ < 0 is not physically meaningful for our system. Thus the equilibrium point eventually becomes arbitrarily far from *P*_0_ in finite time. A graph of the location of *P*_1_ as a function of various values of *b* is given in figure 1.4. We see that as *b* → 0, the equilibrium point diverges with *z*_1_ → −∞.

**Fig. 1.4.**
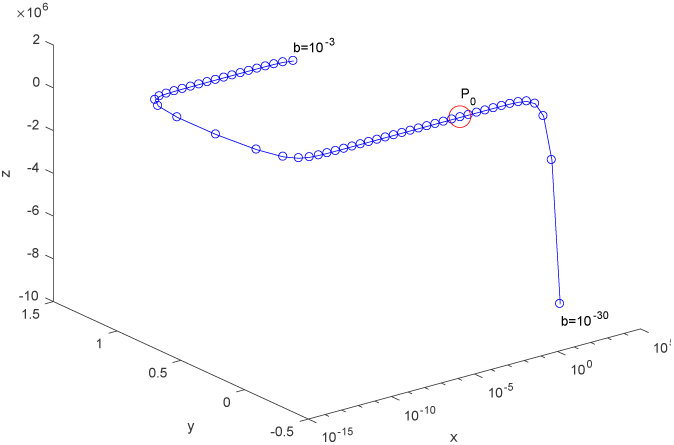
Position of *P*_1_ = (*x*_1_, *y*_1_, *z*_1_) as a function of *b* in blue. *b* ranges from 10^−30^ to 10^−3^. Parameters are: *λ* = 10^−11^, *d* = 10^−9^, *a* = 10^−12^, *β* = 10^2^, *n* = 5. Individual dots for *b* are logarithmic increments of 10^0.5^. We see that *z*_1_ → −∞ as *b* → 0. Location of *P*_0_ is highlighted in red. *P*_0_ and *P*_1_ never coincide precisely, despite appearances using this axis scaling.

Figure 1.5 shows the system asymptotically approaching equilibrium point *P*_1_ as *t* → ∞. We find that *P*_1_ is asymptotically stable for this representative choice of parameters and where *λ* > 0.

**Fig. 1.5.**
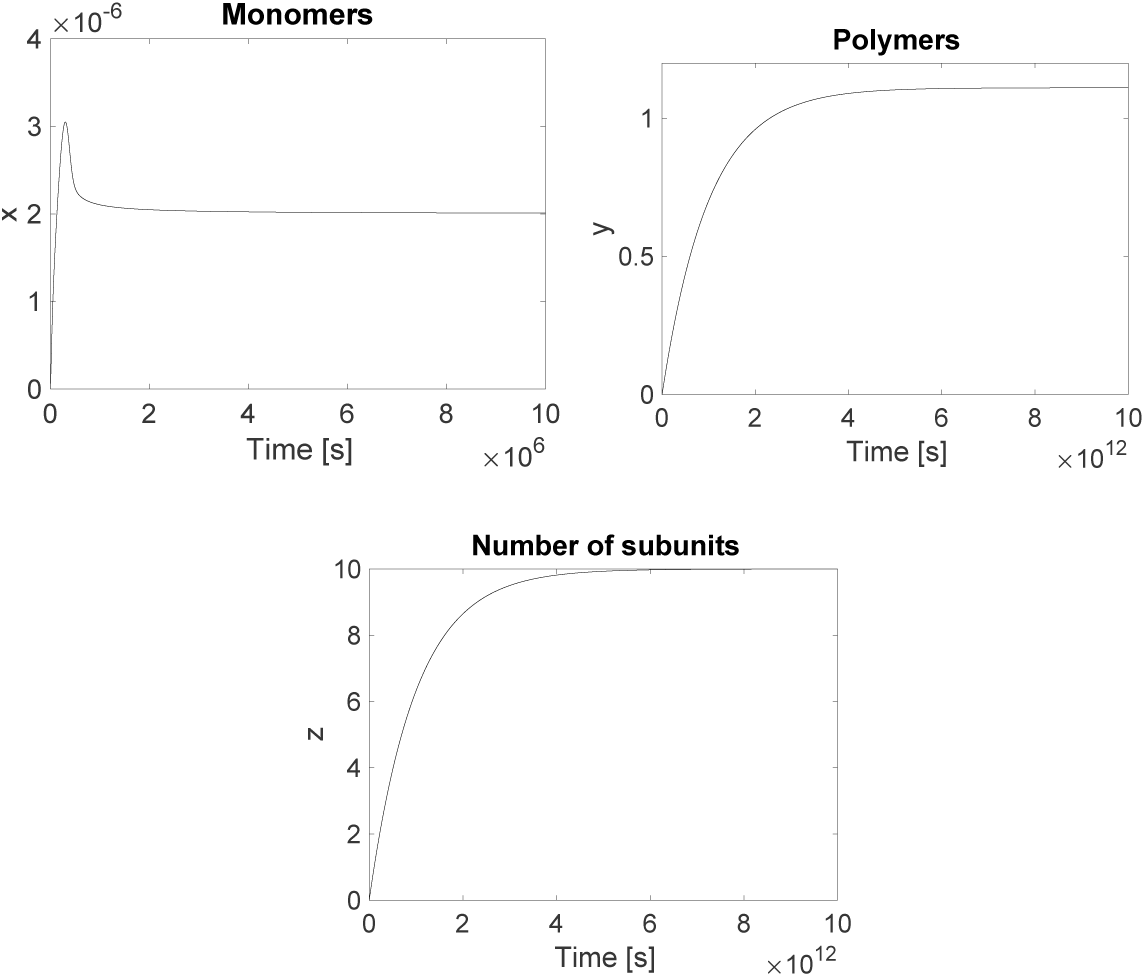
Monomers *x*(*t*), polymers *y*(*t*), and density of subunits *z*(*t*) as functions of time. Parameters were: *λ* = 10^−11^, *d* = 10^−9^, *a* = 10^−12^, *β* = 10^2^, *n* = 5, *b* = 10^−5^. Initial condition is taken as (0, 3.5714×10^−8^, 10^−6^). System asymptotically approaches *P*_1_ ≈ (2.0000 × 10^−6^, 1.1109, 9.9980).

In order to test whether the system approaches *P*_1_ for other values of *b*, we next perform a sensitivity analysis on the fragmentation rate to see how the steady-state solution varies with *b*. Figure 1.6a shows equations (1.1) - (1.3) solved numerically for various values of *b*. Note that in this figure we have taken *λ* > 0 which represents an essentially unlimited supply of monomers. The figure shows that *z*(*t*) goes through a transient decrease period for some values of *b*, which is more pronounced for larger values of *b*. After this, all trajectories eventually increase roughly exponentially for some time. Figure 1.6b is the same as 1.6a but over a longer timeframe, and only for select values of *b*. Here we see that when *b* = 10^−6^, *z*(*t*) approaches *z*_1_ ≈ 9.9998, the *z*-coordinate of equilibrium point *P*_1_. (The same behavior is true for other values of *b* > 0.) This suggests that trajectories with *b* > 0 approach *P*_1_ asymptotically. However, for *b* = 0, *P*_1_ does not exist; the trajectory eventually collapses.

**Fig. 1.6.**
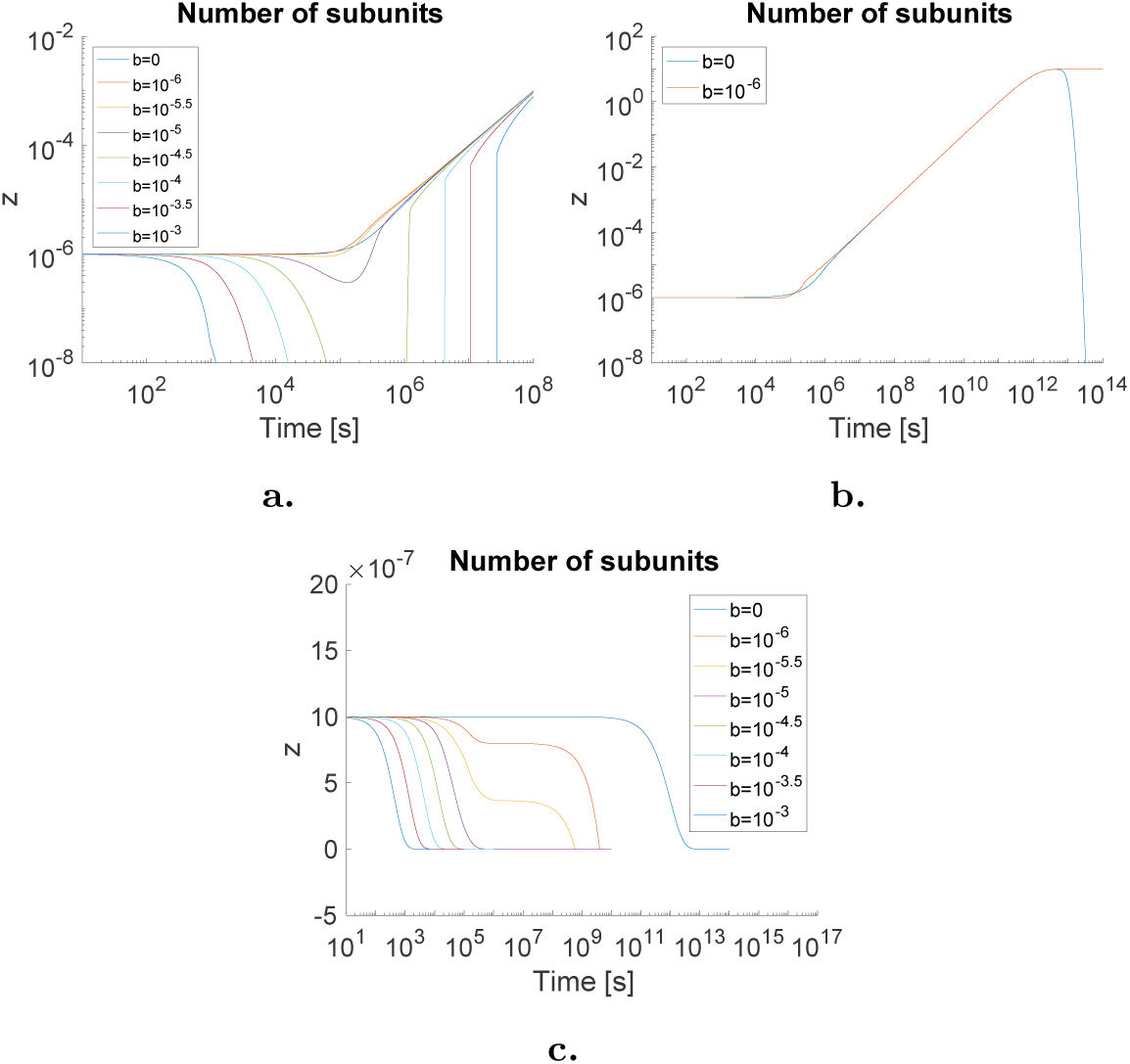
Sensitivity analysis of *z*(*t*) for various values of *b* and *λ*. Subfigures **a** and **b** show *λ* = 10^−11^, and subfigure **c** shows *λ* = 0. In subfigure **a** we see that *z*(*t*) goes through a transient decrease period for some values of *b*, which is more pronounced for larger values of *b*; all trajectories eventually increase roughly exponentially. Sub-figure **b** is a zoom-out of subfigure **a** for select representative values of *b*. Subfigure **b** shows the trajectory for *b* = 10^−6^ approaching *z*_1_ while the trajectory for *b* = 0 eventually collapses. Note that 9.8 ≲ *z*_1_ ≲ 9.9998 when 10^−6^ ≤ *b* ≤ 10^−3^ whereas *P*_1_ does not exist for *b* = 0. In subfigure **c**, all trajectories asymptotically approach *P*_0_ regardless of *b*. All other parameter values and initial conditions are identical to those given in figure 1.5.

Figure 1.6c again shows equations (1.1) - (1.3) solved numerically for various values of *b*, but now with *λ* = 0. This represents a situation where the number of monomers is constrained. In this case, all trajectories asymptotically approach *P*_0_ = (0, 0, 0). This implies that the presence of free monomers is necessary for prion growth.

To understand the evolution of the polymers themselves (i.e. of the distributions *y*_*i*_ over time) we must bound their maximum possible length in equations (1.1) - (1.3) for the purpose of completing numerical simulations. We denote by N the upper bound on the length of polymers, with N sufficiently large (*N* ≫ *n*). We assume *y*_*i*_ = 0 for all *i* > *N*. After some algebraic manipulations, the finite dimension fragmentation dynamical systems become:

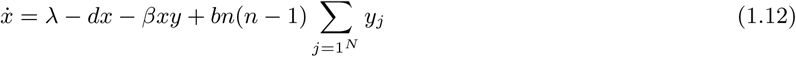

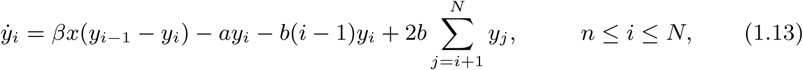

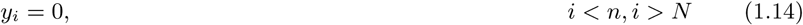

Clearly system (1.12) - (1.14) tends to system (1.1) - (1.3) as *N* → ∞. Figure 1.7a shows the distribution of a Gaussian distribution for a given set of parameters over a short time frame. Note that due to the assumptions of the fragmentation model, polymers of length *i* less than *n* = 5 immediately disintegrate and thus have *y*_*i*_ = 0. Overall, longer polymers gradually defragment to polymers of shorter lengths, until the highest concentration is polymers of length *n* = 5. Figure 1.7b shows the same distribution, but over a much longer time scale; as this set of initial conditions asymptotically approaches *P*_1_, we see a very slow but gradual growth of polymers of all lengths. This very slow growth is representative of prion diseases which have an incubation period of many years after infection before symptoms appear. Figure 1.7c shows the evolution of individual polymer density without fragmentation. Evidently the polymers only grow longer, contradicting experimental observation.

**Fig. 1.7.**
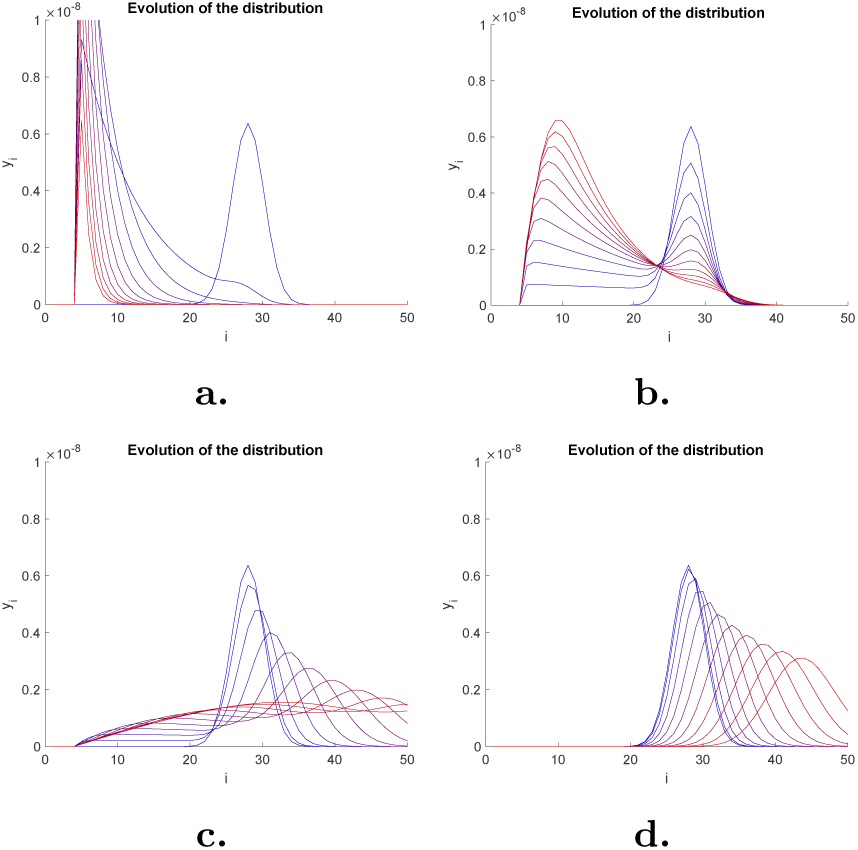
Evolution of a Gaussian distribution initial condition for concentrations of polymers of various lengths, centered at length 28, standard deviation 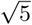, truncated outside of 5 ≤ *i* ≤ 200. Time indicated by color gradient where blue represents *t* = 0 and red represents final time. Subfigure **a** has *b* = 10^−3^ and final time *t* = 10^3^. Subfigure **b** has *b* = 10^−6^ and final time *t* = 10^5^. Subfigure **c** has *b* = 10^−7^ and final time *t* = 3 × 10^5^. Subfigure **d** has *b* = 0 and final time *t* = 2 × 10^5^. Other parameter values were taken as *λ* = 10^−11^, *d* = 10^−9^, *a* = 10^−12^, *β* = 10^2^.

### 1.3 SuPrP Model

The fragmentation model described above has been proposed to be responsible for the multiplication of the templating interface. Recent results demonstrated the existence of a constitutional detailed balance between PrP^Sc^ assemblies and their elementary brick suPrP [10]. The fact that suPrP harbors all the replicative and strain structural determinant could intrinsically constitute a way to increase non-linearly the number of templating interface during the prion replication.

The primary difference with the prior model described in Section 1.2 is the disappearance of a natural fragmentation process. It is replaced by the dynamic interaction of polymerization and depolymerization of the bricks. In [10], the authors study the intimate architecture of infectious prion assemblies through a sequential unfolding and refolding process of the abnormal conformer PrP^Sc^. As explained in the introduction, the unfolding process shows the existence of small oligomeric units, designated as suPrP. In their isolated status they are innocuous but become infectious during the refolding process when stacked into larger assemblies.

Based on the experimental data, suPrP are assumed to be in the range of a PrP dimer or trimer which will be one of the main assumptions in our model. Another key assumption of our model is that alone the suPrP is PK sensitive with very low templating activity, but that once polymerized it becomes a PK-resistant assembly with templating and replicative activity. Finally, we assume that cohesion of PrP monomers in suPrP oligomers involves a strong interaction (which results in suPrP not disassembling back into PrP^C^) while the stacking of suPrP is done through a weak interaction allowing the detailed balance between PrP^Sc^ assemblies and suPrP.

#### Notation

In the new model, *x* still represents the concentration of monomers as in the fragmentation model. We denote the concentration of isolated suPrP bricks as *y*_1_ and the concentration of polymers of length *i* as *y*_*i*_, for *i* ≥ 2 (still called PrP^Sc^, but now assumed to be composed of suPrP).

#### Assumption

We assume the suPrP are trimers, so that each brick is made up of three monomers. The degradation rates *d* and *a* still apply to monomers and polymers, respectively, but since suPrP is highly stable, we assume no degradation of *y*_1_. We make the additional assumption that monomers do not form isolated suPrP (they form the bricks when attaching to polymers), although we will explore in forthcoming work the possibility that this does occur, along with a reverse reaction.

We introduce the coefficients *β* to account for the PrP trimer polymerization with polymers and *p* to account for suPrP polymerization. Then, 3*βxy*_*i*_ represents the rate at which monomers attach to a polymer of length *i* and *py*_1_*y*_*i*_ is the rate at which suPrP attaches to a polymer of length *i*. The rate of depolymerization for polymers of length *i* > 1 is given by *ky*_*i*_ where *k* is the depolymerization coefficient. Based on our assumptions, and contrary to the fragmentation model, there is no *critical size* below which PrP^Sc^ is unstable. Finally, as before, *λ* represents the production of monomers. The interplay between the polymerization and depolymerization processes is depicted in Figure 1.8.

**Fig. 1.8.**
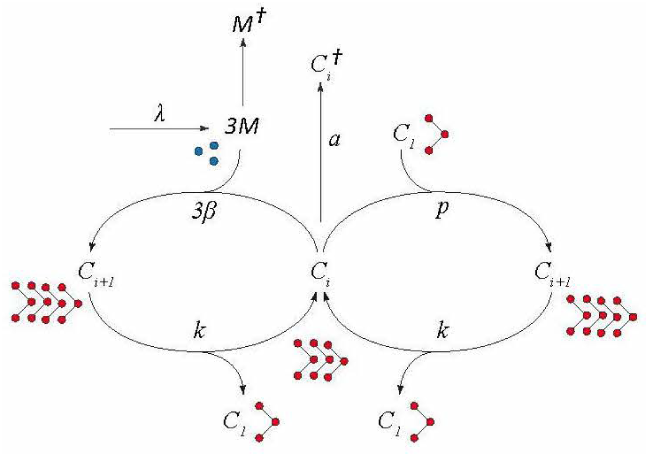
Canonical kinetic model of prion replication including the detailed balance between PrP^Sc^ and suPrP.

The dynamical system is now expressed by the following equations:

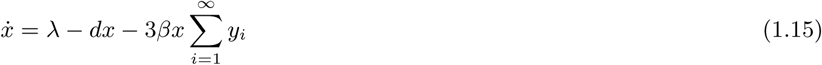

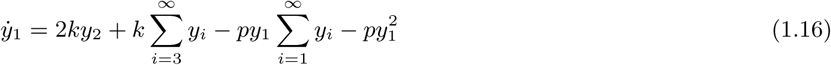

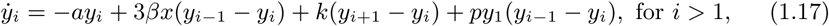

Again, the *dx* term in equation (1.15) should be read as the rate constant *d* times the population of monomers *x*, not as an infinitesimal.

From equation (1.15) we deduce that the concentration of monomers can increase only through the production rate *λ* and no longer through fragmentation. In particular if the production is zero the concentration of monomers will reach zero in finite time. Using as before the notation *y* = ∑*y*_*i*_, the system above is equivalent to:

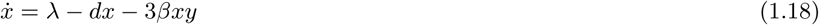

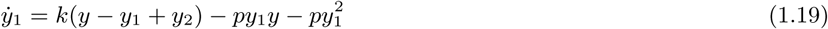

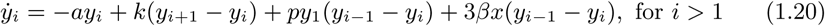

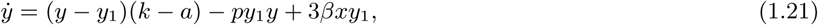

which represents an infinite dimensional model. For the purpose of numerical simulation, we reduce the system to a finite-dimensional one. Since *y* < ∞ and *y*_*i*_ ≥ 0 for all *i*, we have lim_*i*→∞_ *y*_*i*_ = 0. We truncate all values of *i* larger than some threshold value which we call *N*. As long as *N* is taken sufficiently large, this will not impact the dynamics of the system. We thus arrive at the final form of our model:

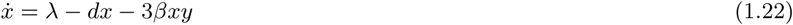

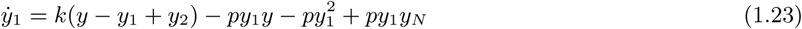

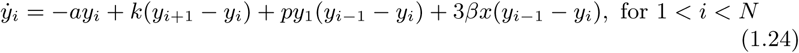

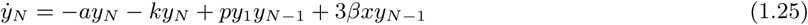

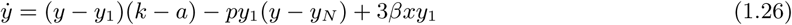

Clearly the model must be such that with the absence of monomers and death the number of bricks must remain constant. We have the following lemma.

##### Lemma 1.

*Assume there is no monomers: x* ≡ 0, *and that a* = 0. *Then, for systems (1.19)-(1.21) and (1.23)-(1.26) we have that* 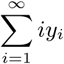 *is constant.*

*Proof.* It follows from a straight calculation that 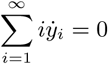.

As in the fragmentation model, we reintroduce the notation 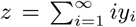 for the rest of the paper since this value is an invariant for our solutions. *z* represents the assembly concentration, but is now measured in suPrP brick-equivalents (not monomers, as we had in the fragmentation model). This means that since *z* is constant our system evolves on an hyperplane in the state space ℝ^*N*^ when *x* ≡ 0, and the constant *z* is given by the initial distribution of *y*_*i*_’s.

#### 1.3.1 Equilibria

We begin by analytically investigating the equilibria for our new model given by equations (1.22) - (1.26). For mathematical simplicity, we consider the case where *λ* = 0. Recall that *λ* represents the formation of monomers by the cell and thus *λ* = 0 could occur in an *in vitro* sample where the initial concentration of PrP^C^ is fixed. We also assume that the death rate *a* = 0 for simplicity since it is usually negligible compared to the other terms.

##### Lemma 2.

*When λ* = *a* = 0, *the equilibrium points of system* (1.22) − (1.26) *are the origin and the one-parameter family given by:*

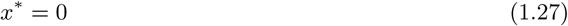

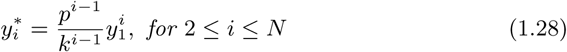

*with y*_1_ *is the free parameter.*

*Proof.* From lemma 1 we have 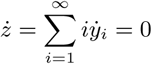, therefore for the equilibrium we have *N* − 1 equations in *N* unknowns and the equilibrium points form a one dimensional curve in ℝ^*N*^ parametrized by the number of individual suPrP bricks *y*_1_. Indeed, from equation (1.25) we deduce that 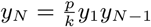. Inserting into equation (1.24) the terms *py*_1_ *y*_1_ cancel out and we have the relation 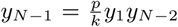. Applying the argument iteratively we obtain 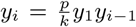. In particular for *i* = 2 we have 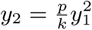. The other terms are obtained by replacing into the recursive formula.

We remark that if we do not assume *a* = 0, the recursion formula becomes 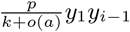 and the calculations can be carried out the same way.

##### Lemma 3.

*When λ* = 0, *the equilibrium points of system* (1.18) − (1.21) *are the origin and* 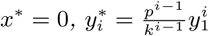 *for all i* ≥ 2.

*Proof.* It is a straightforward calculation that all the derivatives vanish.

From lemma 2, it can be seen that the equilibrium distribution depends on the ratio 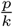. In particular if we normalize the distribution by *y*_1_, we have that

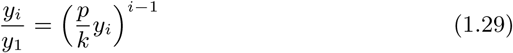

which decays exponentially with respect to the length of the polymers and with respect to the ratio *p/k*. Moreover, we have that for a given distribution of *y*_*i*_:

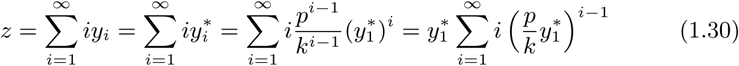

where 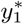 is the amount of suPrP bricks when the distribution relaxes to the equilibrium. Using 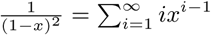 for |*x*| < 1 we have:

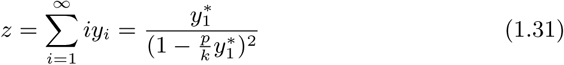

Recall that the value (1.30) is an invariant for the solutions (provided that there are no monomers and *a* = 0). In particular, from formula (1.29) the equilibrium is completely determined once the number of suPrP bricks 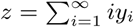 is fixed, and it is given by formula (1.28) with:

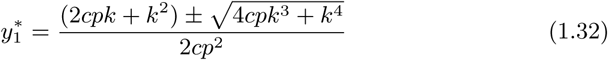

Although there are two solutions here, choosing the + results in a value of 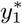 that violates the convergence of the series 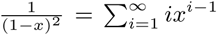. So, are left with only the − solution:

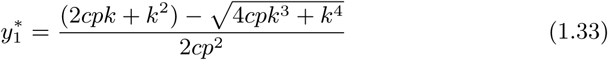

The moment of order two at equilibrium is given by 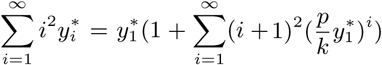. This series does not have a closed form formula as in (1.29), hence distribution with the same moment of order two at equilibrium might reach different equilibria.

#### 1.3.2 Polymer Evolution

The primary goal of our simulations is to demonstrate the interplay between the suPrP bricks and the polymers of higher length. It will show that fragmentation is not the key process between the dynamics of prion replication. We divide the numerical simulations into two, one corresponding to continuous production of monomers while the other one has limited monomer resources which decrease to zero in finite time.

##### Case I *λ* > 0

This corresponds to having an unlimited number of monomers. Evolution of the polymers is depicted in Figure 1.9. We see that with *λ* positive and sufficiently large, the concentration of small polymers rises and quickly dominates the initial distribution.

**Fig. 1.9.**
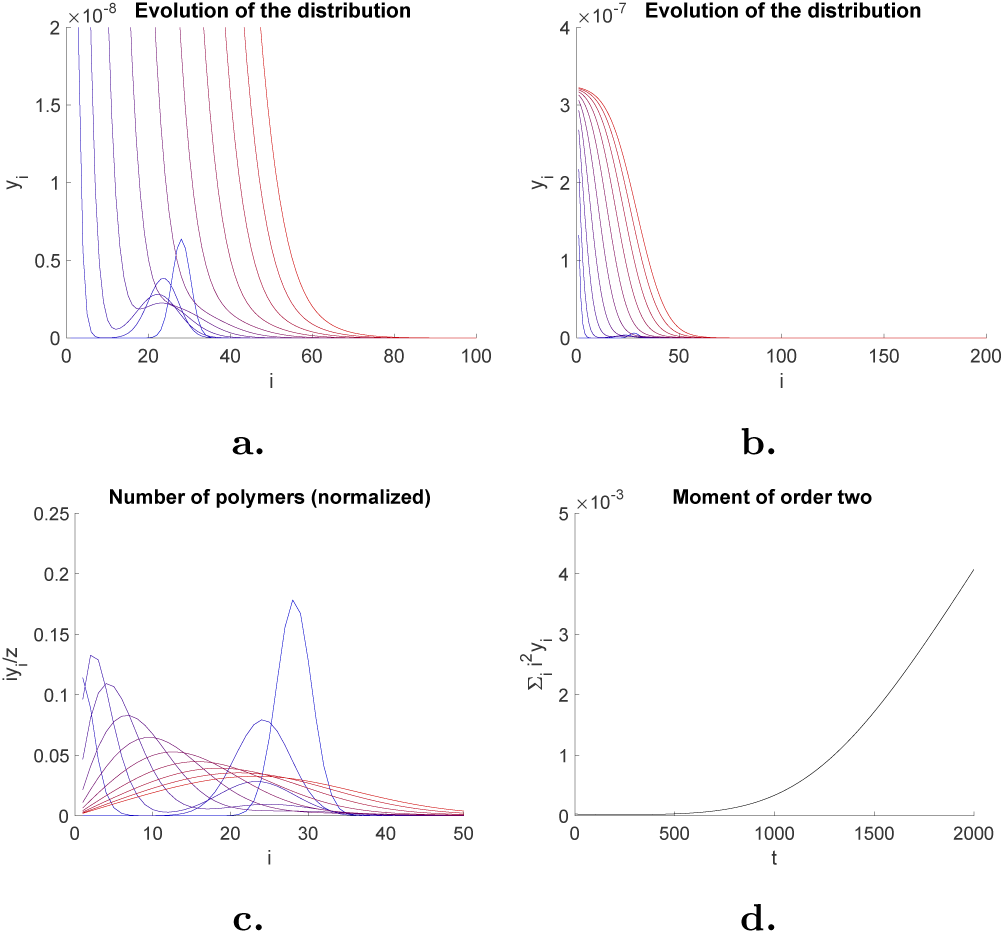
Subfigures **a** and **b** show evolution of the distribution *y*_*i*_ from the initial distribution at *t* = 0 (blue) to the final distribution at *t* = 2 × 10^3^ (red); these are the same figure with different vertical axis scalings. Subfigure **c** shows the number of polymers normalized by the assembly concentration (suPrP brick-equivalents) ∑*iy*_*i*_*/z*, and subfigure **d** shows moment of order two ∑*i*^2^*y*_*i*_ as functions of time. Parameters are: *λ* = 10^−7^, *d* = 10^−9^, *a* = 10^−12^, *β* = 10^2^, *k* = 1*/*3 × 10^−1^, *p* = 10^5^, *N* = 200. Initial conditi*v*o<u>n</u> for *y*_*i*_ was a truncated Gaussian distribution centered at 28, standard deviation 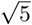. Initial condition for *x* was *x* = 0.

##### Case II

*λ* = 0 This corresponds to an initial finite quantity of monomers that decreases to zero since there is no production and suPrP bricks do not degrade. As a result, in finite time the system will reach an equilibrium depending on the rate constant values. For simplicity, we also set *x* = 0 for the initial condition which implies *x* ≡ 0 throughout the evolution. It can be seen on figure 1.10 that the initial polymer distribution collapses downward and to the left while the concentration of the smallest polymers increases rapidly. Qualitatively this largely matches the behavior we saw in the fragmentation model of figure 1.7. Thus this represents a proof-of-concept that the existence of the bricks will result in a similar outcome for the population dynamics over time to that of fragmentation.

**Fig. 1.10.**
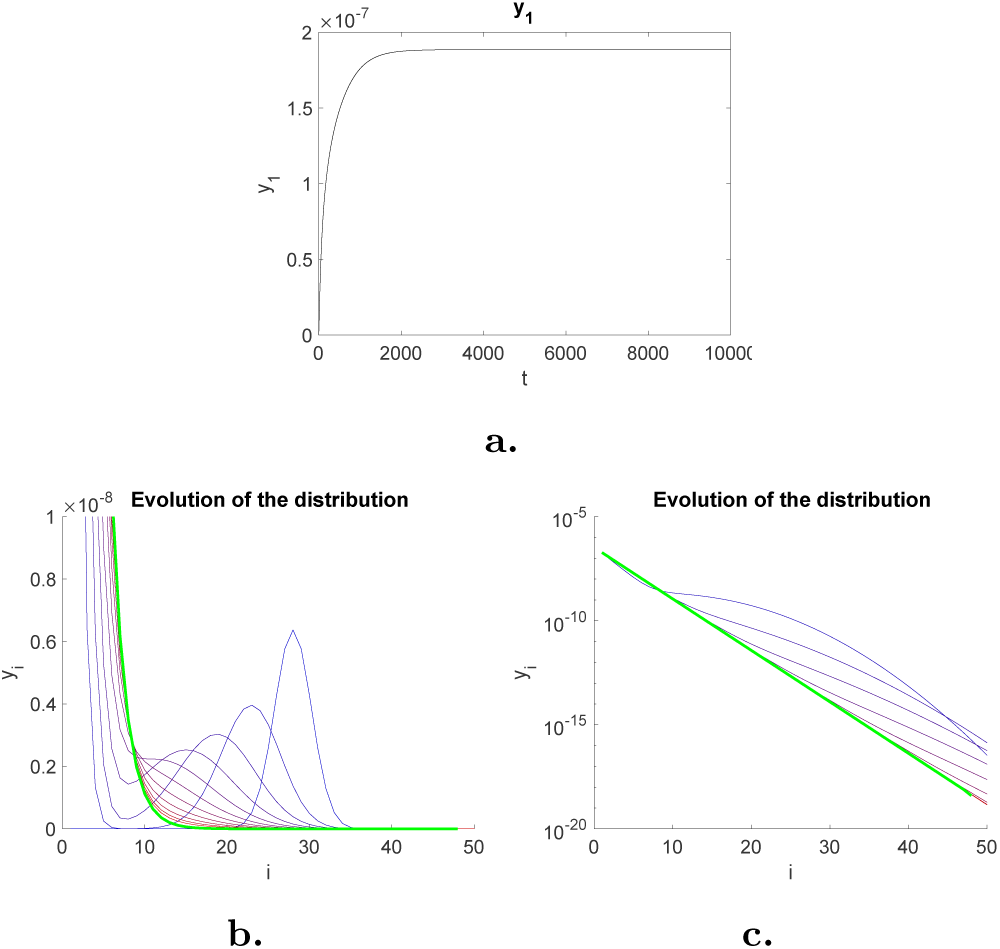
Subfigure **a** shows that *y*_1_ asymptotically approaches a value of ≈ 1.8858 × 10^−7^ as *t* → ∞. Subfigure **b** shows evolution of the distribution *y*_*i*_ from the initial distribution at *t* = 0 (blue) to the final distribution at *t* = 2 × 10^3^ (red). Green distribution represents the equilibrium point of equations (1.27) - (1.28), taking *y*_1_ to be the numerical value shown in subfigure **a**. Subfigure **c** also shows the same evolution but over a longer time scale with blue being time *t* = 10^3^ and red being final time *t* = 10^4^, and with logarithmic vertical axis scaling. The evolution asymptotically approaches the equilibrium point. Other parameters were: *λ* = 0, *d* = 10^−9^, *a* = 0, *β* = 10^2^, *k* = 1*/*3 × 10^−1^, *p* = 10^5^, *N* = 200. Initial condition for *y*_*i*_ was a truncated Gaussian distribution centered at 28, standard deviation 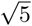. Initial condition for *x* was *x* = 0.

In subfigure 1.10a we see that *y*_1_ approaches a value of ≈ 1.8858×10^−7^ as *t* → ∞. This is a numerical validation of equation (1.33) with our choice of parameters *p* = 10^5^, *k* = 1*/*3 × 10^−1^, and *z* = 10^−6^. This value was then used to generate the equilibrium point found analytically in equations (1.27) - (1.28). Subfigures 1.10b and c show the evolution of the distribution with initial distribution in blue and final distribution in red. These figures have linear and logarithmic vertical scalings, respectively. In both cases we see the asymptotic approach of the evolution to the equilibrium point in green. This is a numerical validation of our analytical solution for the equilibrium point of equations (1.27) - (1.28).

To investigate the dependence of 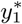 on the choice of parameters, in figure 1.11a we show 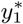 as a function of time for several simulations where *p* was varied. Each asymptotically approaches a different value of 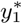 as *t* → ∞. Figure 1.11b shows 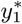 as a function of *p* from equation (1.33). We see numerical verification that the equilibrium values of 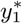 do indeed approach the values found analytically in equation (1.33).

**Fig. 1.11.**
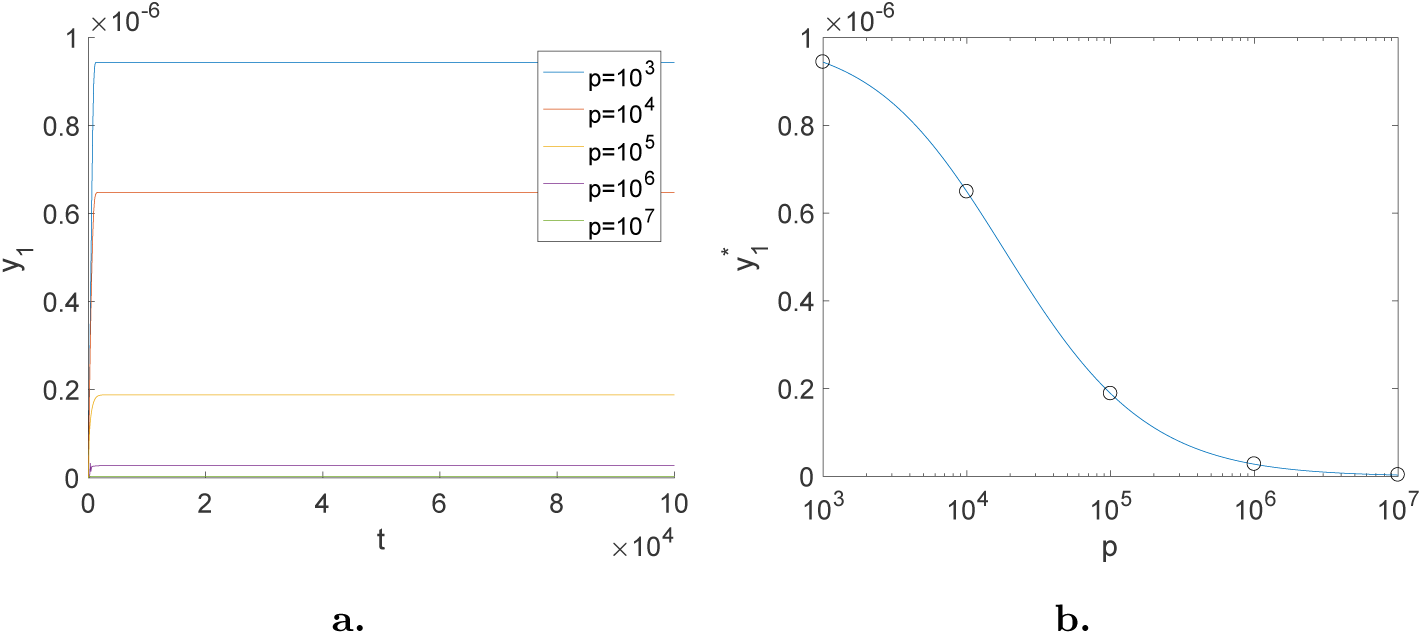
Subfigure **a** shows a superposition of 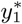 as a function of time for different values of *p. p* values were between 10^3^ and 10^7^ as labeled. Initial condition for *y*_*I*_ was a truncated Gaussian distribution centered at 28, standard deviation 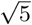. Initial condition for *x* was *x* = 0. Other parameters were: *λ* = 0, *d* = 10^−9^, *a* = 0, *β* = 10^2^, *k* = 1*/*3 × 10^−1^, *N* = 200. *z* = ∑*iy*_*i*_ was fixed at 10^−6^. Subfigure **b** shows 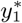 as a function of *p* from equation (1.33). Black circles indicate numerical values of 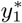 at *t* = 10^5^ from subfigure **a**. This is a numerical verification that our analytical equilibrium calculation of equation (1.33) is correct.

The role of the parameters *k* and *p* is illustrated in figure 1.12. Three values of the ratio *p/k* are shown. Recall that parameter *p* represents the rate of polymerization while parameter *k* represents depolymerization. The ratio *p/k* indicates the relative speed of these two processes. *p/k* small indicates more rapid depolymerization, which we see illustrated in figure 1.12b, the number of polymers, normalized by the assembly concentration (suPrP brick-equivalents), with rapidly increasing levels of small *i*. On the other hand, *p/k* large indicates more rapid polymerization; see figure 1.12k where the initial hump spreads, but we do not see a spike for small *i*.

**Fig. 1.12.**
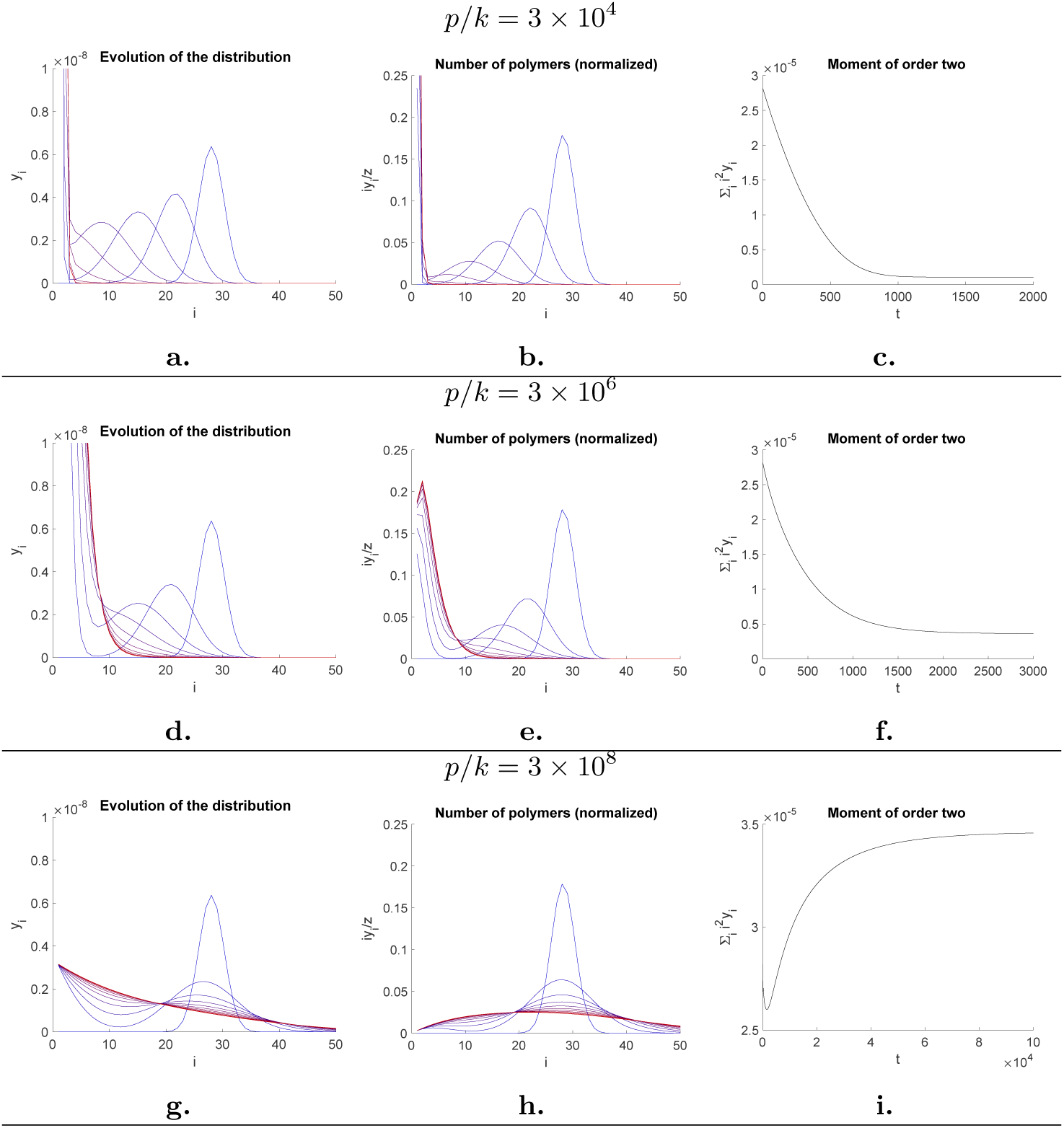
Sensitivity analysis on parameters *k* and *p*. Subfigures **a,d,g** show evolution of the distribution *y*_*i*_. Subfigures **b,e,h** show the number of polymers, normalized by the assembly concentration (suPrP brick-equivalents) *iy*_*i*_*/z*. Subfigures **c,f,i** show moment of order two ∑*i*^2^*y*_*i*_ as functions of time. For subfigures **a,b,c**, *p* = 10^3^, for *p/k* = 3 × 10^4^, and final time *t* = 2 × 10^3^. For subfigures **d,e,f**, *p* = 10^5^, for *p/k* = 3 × 10^6^, and final time *t* = 3 × 10^3^. For subfigures **g,h,i**, *p* = 10^7^, for *p/k* = 3 × 10^8^ and final time *t* = 5 × 10^3^ (except subfigure **i**, which has final time *t* = 10^5^). For all subfigures, other parameters were: *λ* = 0, *k* = 1*/*3× 10^−1^, *d* = 10^−9^, *a* = 10^−12^, *β* = 10^2^, *N* = 200. Initial condition for *y*_*i*_ was a truncated Gaussian distribution centered at 28, standard deviation 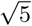. Initial condition for *x* was *x* = 0. Blue is initial distribution and red is final time, with color gradient in between.

The moment of second order is a direct experimental measurement of the weighted-mean average size distribution. It is directly determined from light scattered intensity of the sample containing protein assemblies using Rayleigh relation [1]. As shown in figure 1.12 for *p/k* = 3.106 the second order momentum initially decrease to reach a plateau whereas for *p/k* = 3.108 it initially decreases and then increases to a plateau which is due to a dissipation phenomenon.

In figure 1.13, we conduct a sensitivity analysis on the initial distribution. We take the initial condition to be a truncated Gaussian distribution centered at *µ* with standard deviation 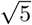. *µ* was varied between 20 and 100. Again, the green timeframe highlights the time at which the moment of order two is minimum.

**Fig. 1.13.**
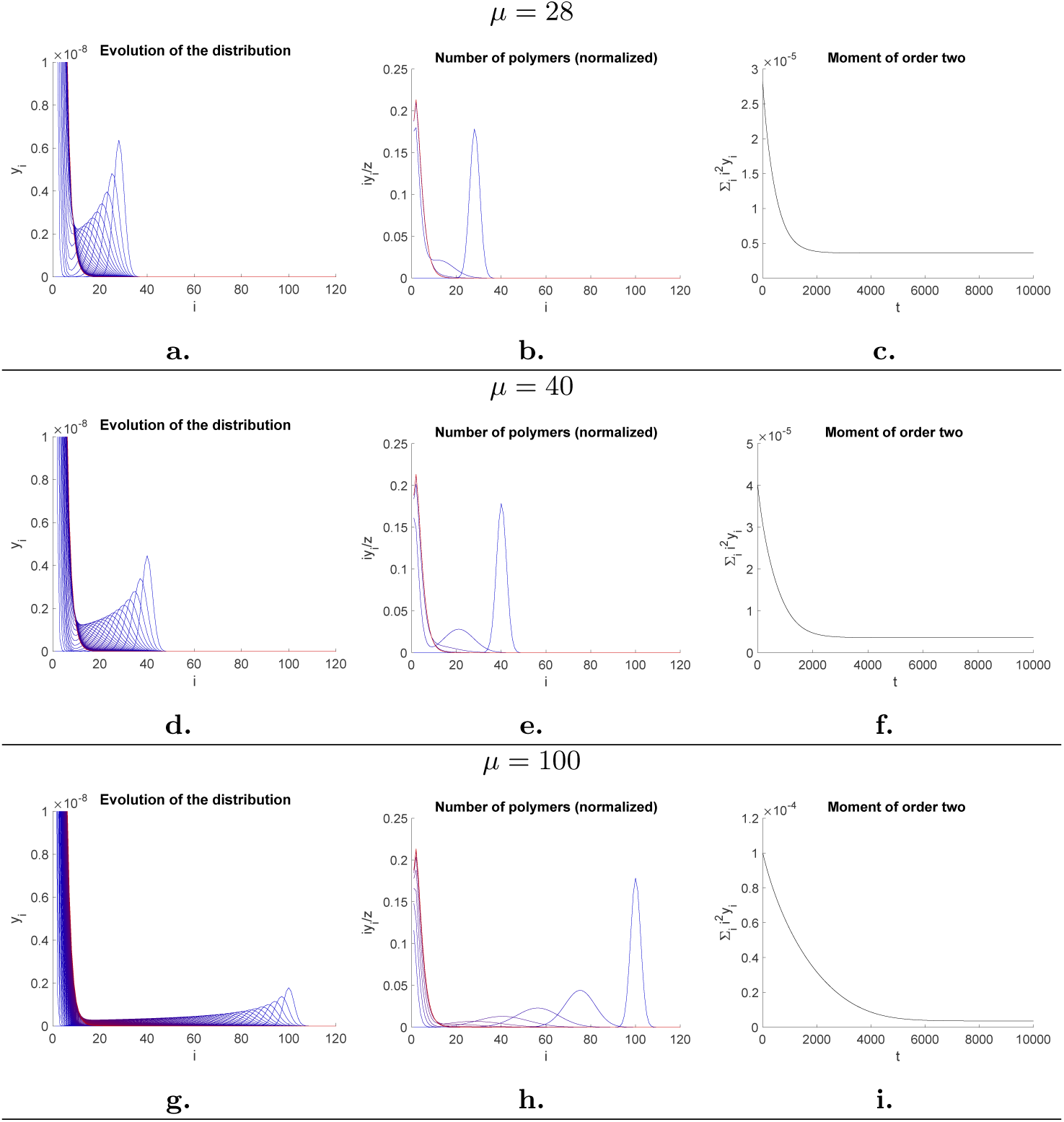
Sensitivity analysis on mean *µ* of truncated Gaussian initial distribution for *y*_*i*_. Subfigures **a,d,g** show evolution of the distribution *y*_*i*_. Subfigures **b,e,h** show the number of polymers, normalized by the assembly concentration (suPrP brick-equivalents) *iy*_*i*_*/z*. Subfigures **c,f,i** show moment of order two ∑*i*^2^*y*_*i*_ as functions of time. For subfigures **a,b,c**, *µ* = 28. For subfigures **a,b,c**, *µ* = 40. For subfigures **a,b,c**, *µ* = 100. For all subfigures, other parameters were: *λ* = 0, *k* = 1*/*3 × 10^−1^, *p* = 10^5^, *d* = 10^−9^, *a* = 10^−12^, *β* = 10^2^, *N* = 200. Initial condition for *x* was *x* = 0. Blue is initial distribution and red is final *t* = 1×10^4^, with color gradient in between.

#### 1.3.3 Discussion

The prion replication can mechanistically be divided into two entangled steps. The first step corresponds to the templating process where the mechanisms is related to Koshlan fitted induced adjustment [11] and corresponds to a linear perpetuation of the structural information. During the templating process the number of interfaces remains constant. The second step corresponds to the amplification of the structural information by the multiplication of the templating interface. In the case of fungus prions, the amplification process has been reported to be related to the fragmentation induced by heat-shock proteins (hsp) [15, 16, 17]. The cytological localization of fungus prions such as Ure2 and sup35 assemblies makes them in close contact with hsp-families and dNTP as source of energy rendering therefore their fragmentations possible as an active and energy consuming process. In mammalian prion disease the amplification of the pathological information is far from clear. In the case of PrP^Sc^ assemblies, the cytological localization of the replication process makes the encounter of PrP^Sc^ assemblies with proteins that could act as a fragmentase highly improbable. The existence of constitutional detailed balance between PrP^Sc^ assemblies and their elementary building blocks suPrP constitute an alternative mechanism to circumvent the hypothetical fragmentation process. In this paper, we demonstrated in section 1.3.2 that this new paradigm does produce the necessary amplification phase to multiply the templating interface. It can be seen very clearly through our simulations as the quantity of the suPrP is increasing throughout the evolution of the system.

In particular, we carried out analytical calculations when *x* ≡ 0 (or more generally the case when *λ* = 0 since the system eventually uses up all the monomers and converges to the situation *x* ≡ 0) and *a* = 0 (no destruction of polymers). In that case the assembly concentration (in suPrP-equivalents), given by 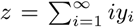, is an invariant of the evolution of the system. For the truncated system (1.23)-(1.26) it means that the system evolves on a hyperplane and we can reduce its dimension to *N* − 1. Once *z* is fixed, the equilibrium is completely determined by the ratio 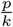 of the polymerization and depolymerization constant rates. Forthcoming work will analyze the stability of the equilibria (there is numerical evidence suggesting that the system is asymptotically stable). When *λ* ≠ 0, the situation is more complicated since we have creation of monomers external to the system. Therefore *z* is not an invariant anymore and the geometry is more intricate (the system does not evolve on an hyperplane).

### 1.4 Prion Size Modulation by Temperature Adjustment

Experimentation has shown through the moment of order two *m*^2^ = ∑*i*^2^*y*_*i*_ that temperature impacts the dynamics of prion assemblies. This is a consequence of the fact that the dependence of the rate constant of a chemical reaction on the absolute temperature is given by the Arrhenius equation. The model presented in Section 1.3 corresponds to fixed reaction coefficients, therefore representing a prescribed temperature. This suggests that temperature can be used as a control parameter to steer the system from an initial equilibrium to a final one.

#### 1.4.1 Optimal Control Problem

In this section we introduce the idea of controlling the rate parameters by introducing them as varying parameters rather than constants. As a first step, we consider time dependence but in forthcoming work the temperature dependence will be introduced as well.

#### Assumption

For simplicity, in this model we assume *x* ≡ 0. The rate constants *λ, d* and *β* are therefore neglected in equations (1.23)- (1.25).

#### Notation

In this model we define the control *u* = (*u*_1_, *u*_2_, *u*_3_)^*t*^ given by the degradation (*a* = *u*_1_), depolymerization (*k* = *u*_2_) and polymerization (*p* = *u*_3_) rates. We assume *u* is a measurable bounded function.

The dynamical system of prion assemblies is now a driftless affine control system given by:

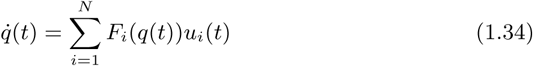

where *q* = (*y*_1_, *y*_2_ … *y*_*N*_)^*t*^. The vector fields *F*_*i*_ are given by equations (1.23) - (1.25) with 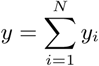. Given a fixed set of rate parameters, we associate its unique nonzero equilibrium *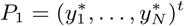* given by Lemma 2. Therefore, we can control stationary states of prion assemblies by varying the control. Moreover, we might want to accomplish the transition between stationary states while minimizing time or some internal energy of the system. In this paper, we focus on time minimization. We can now state the control problem as follows.

#### Control Problem

*Given two equilibria* 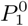 *and* 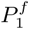, *find a measurable bounded function u*(.) *that steers the prion assemblies from* 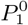 *to* 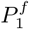 *in minimum time.*

We assume the domain of control to be a parallelogram, i.e. 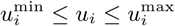 for each *u*_*i*_ with 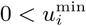 for *i* = 2, 3. This comes from the fact that the rate parameters are governed by the Arrhenius law with coefficients that are parameter dependent. We define the control domain as 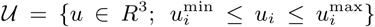 where the bounds are fixed scalars. The control problem is stated in all generalities, and from a practical aspect the control will then have to be approximated by a continuous function (this will be the topic of forthcoming work; here we restrict our study to the analysis of extremals).

In this paper we will restrict ourselves to the case *N* = 3 for an initial analysis of the abnormal extremals. For *N* = 3, we have:

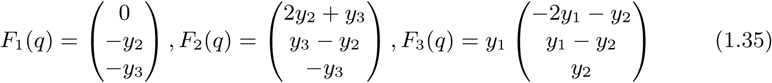

where *y*_1_ is the concentration of SuPrP, *y*_2_, *y*_3_ are respectively the concentrations of polymers of length 2 and 3. The equations can then be written as:

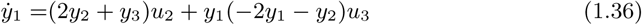

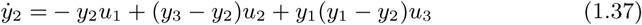

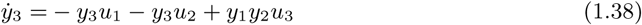

Given 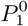 and 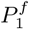, find the time minimal control *u* : [0, *T*] → 𝒰 such that 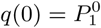 and 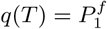.

##### Lemma 4.

*The control vector fields F*_1_, *F*_2_, *F*_3_ *of the affine control system (1.36)- (1.38) are lineraly dependent on*

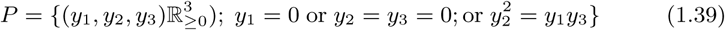

*Proof.* The determinant |*F*_1_, *F*_2_, *F*_3_| is equal to 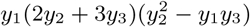.

##### Lemma 5.

*Along a non zero solution we cannot have y*_1_ ≡ 0 *nor y*_3_ ≡ 0.

*Proof.* If *y*_1_ ≡ 0, it implies that 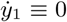 and therefore either *u*_2_ ≡ 0 or *y*_2_ = *y*_3_ ≡ 0. By assumption *u*_2_ > 0 and the origin is an equilibrium point. If *y*_3_ ≡ 0 it implies 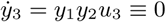 imposing *y*_2_ ≡ 0. Since then 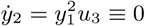 we reach a contradiction.

We remark that the case *u*_1_ = 0 corresponds to a zero death rate for the polymers. In that case the assembly concentration is constant and the system reduces to a 2-dimensional system evolving on an hyperplane defined by *x* + 2*y* + 3*z* =constant. The equilibrium has been computed in a prior section.

#### 1.4.2 Maximum Principle

Assume that there exists an admissible time-optimal control *u* : [0, *T*] → 𝒰, such that the corresponding trajectory *q*(*t*) is a solution of equations (1.36)-(1.38) and steers the prion assemblies from 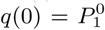 to 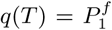. For the minimum time problem, the Maximum Principle, see [6], implies that there exists an absolutely continuous vector *p* : [0, *T*] → *R*^3^, *p* ≠ 0 for all *t*, such that the following conditions hold almost everywhere:

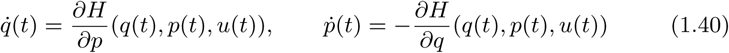

where the Hamiltonian function *H* is given by 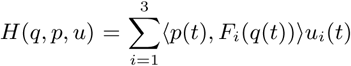.

Furthermore, the maximum condition holds:

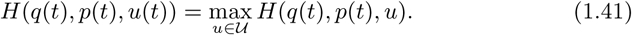

It can be shown that *H*(*q*(*t*), *p*(*t*), *u*(*t*)) is constant along the solutions of (1.40) and is greater or equal to 0. A triple (*q, p, u*) which satisfies the Maximum Principle is called an extremal, and the vector function *p*(.) is called the adjoint vector. A direct consequence of the maximum condition is:

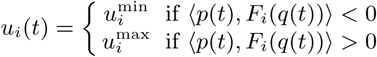

which means that the zeroes of the functions ⟨*p*(*t*), *F*_*i*_(*q*(*t*))⟩ determine the structure of the control. We define:

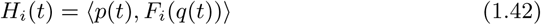

for *i* = 1, 2, 3. A component *u*_*i*_ of the control is said to be *bang-bang* on a given interval [*t*_1_, *t*_2_] *⊂* [0, *T*] if *H*_*i*_(*t*) 0 for almost all *t* ∈ [*t*_1_, *t*_2_]. Therefore, a bang-bang component of the control only takes values in 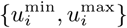 for almost every *t* ∈ [*t*_1_, *t*_2_]. If there is a nontrivial interval [*t*_1_, *t*_2_] such that a switching function is identically zero, the corresponding component of the control is said to be *singular* on that interval. A singular component of the control is said to be *strict* if the other controls are bang-bang. Finally, assume *u*_*i*_ is bang-bang. Then, we say that *t*_*s*_ ∈ [*t*_1_, *t*_2_] is a *i*-switching time if, for each interval of the form]*t*_*s*_ − *ε, t*_*s*_ + *ε*[*∩*[*t*_1_, *t*_2_], *ε* > 0, *u*_*i*_ is not constant.

#### Singular Extremals

Assume the extremal is *u*_*i*_-singular on some nonempty interval [*t*_0_, *t*_1_]. By definition it means that *H*_*i*_(*t*) ≡ 0 on *I*. Differentiating this expression we obtain:

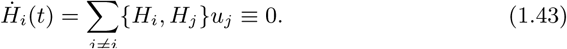

where {,} is the Poisson bracket. We introduce the notation *H*_*ij*_ = {*H*_*i*_, *H*_*j*_} for the rest of the paper.

To understand the singular extremals we clearly need to compute the Lie algebra associated to our vector field since {*H*_*i*_, *H*_*j*_} = ⟨*p*, [*F*_*i*_, *F*_*j*_]⟩.

##### Proposition 1.

*The Lie brackets of order 1 for system (1.36)-(1.38) are given by:*

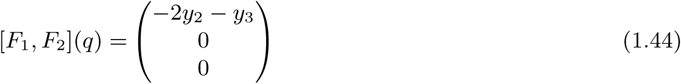

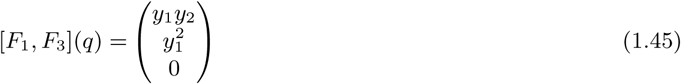

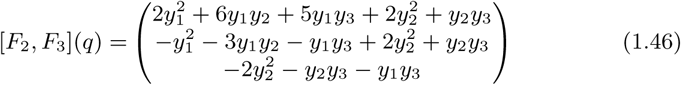

*Proof.* It follows from a straightforward computation.

##### Lemma 6.

*The vector fields F*_1_, *F*_2_, [*F*_1_, *F*_2_] *are linearly independent outside the set y*_3_ = 0. *As a result they form a basis for the set of vector fields.*

*Proof.* The determinant of (*F*_1_, *F*_2_, [*F*_1_, *F*_2_]) is given by 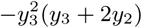.

We say that an extremal is *totally singular* if all three component of the control are singular at the same time. They are characterized by the following properties.

##### Proposition 2.

*A totally singular control in contained in the set*

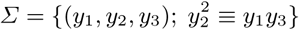

*and satisfies almost everywhere:*

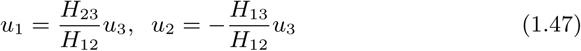

*where u*_3_ *is free.*

*Proof.* This is a direct consequence of the fact that (1.43) must hold for *i* = 1, 2, 3. It can be written as:

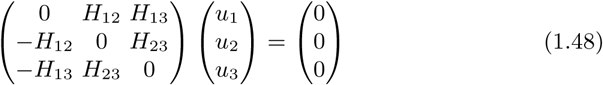

where the 3 × 3 matrix is skew-symmetric and therefore singular. If *H*_12_ ≠ 0 solving for *u*_1_ and *u*_2_ provides the relation between the component of the control. *H*_12_ ≡ 0 is not possible, see lemma 6.

The result above has biological consequences. we recall that we assume the control to be the rates for polymerization and depolymerization as well as the death of polymers and that these vary with respect to the temperature following the Arrhenius law. This means that all controls *u*_*i*_’s should be singular, indeed either the temperature is constant and the constant rate is fixed within its possible bounds or the control is continuously varying. In other words it means that bang extremals are not biologically relevant unless we are at extreme temperatures. Our analysis shows that the set of singular extremals is constrained to satisfy *y*^2^ ≡ *xz*; therefore a more in-depth biological understanding will be necessary before further consideration of optimization. A consequence could be that a system is very limited in terms of which assembly concentration level it can reach by changing the temperature.

## 1.5 Acknowledgments

The authors would like to thank the Simons Foundation for their support. The research presented in this paper is supported by a Simons Collaboration Grants for Mathematicians.

